# Time-gated Fluorescence Lifetime Imaging in the Near Infrared Regime; A Comprehensive Study Toward In Vivo Imaging

**DOI:** 10.1101/2023.05.21.541614

**Authors:** Meital Harel, Uri Arbiv, Rinat Ankri

## Abstract

Fluorescence lifetime imaging has an enormous impact on our understanding of biological systems, both in vitro and in vivo. It is a powerful tool for the non-invasive in vitro and in vivo biomolecular and cellular investigations. In particular, it has the potential to target and multiplex different species with high sensitivity and specificity, providing a fast and noninvasive readout at low cost. In this work, we present a time-saving Monte Carlo (MC) simulation of fluorescent photons scattering within a turbid medium, followed by phasor analyzes which enabled the simple multiplexing of different targets in one frame. We then demonstrate a simple and fast method for wide-field FLI in the near-infrared (NIR) region, where tissue scattering and autofluorescence are significantly lower, to enable imaging of deep tissue, using the state-of-the-art timed single-photon avalanche diode array camera (SPAD), SPAD512S. In particular, we show how phasor scattering increases with depth. However, using appropriate background correction, a simple “cut-off” method, and averaging, we can multiplex two targets in one image to a depth of 1 cm in tissue. Our results show that it is possible to perform in vivo FLI under challenging conditions, using standard NIR fluorophores with short lifetimes.

## Introduction

Fluorescence-based imaging has an enormous impact on our understanding of biological systems, both in vitro and in vivo. It has the potential to target and multiplex different species with high sensitivity and specificity, providing a fast and noninvasive readout at low cost. In vivo fluorescence imaging (FI) enables to study biomolecules and cells behavior in their native environment. These advantages have been showcased in many biomedical investigations, including probing tissue physiology^1^, detecting early stages of disease^2^, sensing molecular concentrations of delivered pharmaceuticals^3^, observing intracellular fluorescent proteins^4, 5^, and much more. However, disadvantages of current in vivo fluorescence imaging include the low penetration of visible light, which is the electromagnetic regime in which most fluorophores emit their fluorescence, through tissue and background autofluorescence. These can be overcome using near-infrared (NIR, or longer) light in biological research or clinical models. Owing to advances in reducing photon scattering, light absorption and autofluorescence, NIR fluorescence affords high imaging resolution with increasing tissue penetration depths.

Live animals imaging requires wide-field (parallel) data acquisition, as it minimizes the temporal delay between data recorded in different regions of the field of view (FOV). It also has the advantage of dispensing with costly and complex scanning devices, even though it creates other challenges. Single photon avalanche diodes (SPAD) cameras capabilities for fluorescence lifetime imaging (FLI) microscopy (FLIM) applications in the visible, as well as the NIR spectrum, have been widely described^6-8^. SPAD cameras implements a fast time-gating control and are capable of measuring not only the fluctuations of the fluorescence intensity at very high frame-rates (50 to 100 kHz)^9, 10^ but also the decay kinetics of fluorophores after illumination by a pulsed laser^10, 11^. The measurement of fluorescence decay kinetics by each SPAD allows the monitoring of ultra-fast photophysical processes, as Förster resonance energy transfer (FRET)^12^, and to identify multiple fluorescence sources. Although studies demonstrated SPAD cameras’ potential for microscopic biological applications, there are still two main challenges involved with using it for small animal NIR imaging applications: (i) near-infrared (NIR) dyes are challenging to detect, due to both the low quantum efficiency of these dyes and the reduced photon detection probability of silicon SPADs in the NIR^6^, and (ii) most NIR dyes exhibit similar, very short lifetimes, of few hundreds of picoseconds (ps)^13, 14^, which make their detection challenging.

In this work, we aim to investigate the performance of timed SPAD arrays in wide-angle NIR FLI. Imaging FLT instead of fluorescence intensity for multiplexing purposes leads to a much less complex optical setup, as different species can be detected in one image with a single excitation wavelength and a single detector. Nevertheless, available NIR dyes have very short lifetimes of up to 1 ns. With such short lifetimes, it is very difficult to distinguish between multiple NIR labels and intrinsic signals. We therefore present a comprehensive study of the multiplexing capabilities of timed FLI. We present FLI simulations and experiments with a single fluorophore as well as with two adjacent fluorophores behind scattering media that mimic skin tissue with increasing thickness. We first investigate this using Monte Carlo simulations (MC) of the path of fluorescent photons in the scattering medium. The MC simulations are a well-known method that allows an in-depth study of the photon path in a turbid medium such as tissue ^15-17^, while recent work has extended the MC model to simulate the path of fluorescent photons in biological tissue, reflecting the growing interest in fluorescence-based imaging for in vivo biomedical applications^18^. For example, Welch et al. proposed the most accurate fluorescence Monte Carlo method (sfMC)^18^, in which the probabilities of emission and absorption of a photon from a fluorophore are calculated in a separate MC code and later combined by convolution. This method is very time consuming, although shorter than the direct approach. In this work, we use a recently proposed simplified MC model for fluorescence photon propagation in biological tissue, in which we neglect the excitation photons and consider the fluorophore as an isotropic light source^19^ (thus avoiding the need for the “method of images”^20^). Despite these neglections, which reduce the simulation time by half, we show that the general behavior of simulation and experiments correlates well. We then use the results of our simulation method in experimental FLT analyzes to provide a fast and accurate method for in vivo FLI simulations.

FLI experiments were performed using SPAD512S^21^, the most advanced large-array camera SPAD, which features a low dark count rate and fully configurable, high-resolution time-gating capabilities that enable fluorescence lifetime imaging with picosecond time resolution and an acquisition time of only 2.6 ms per gate. The resulting data were analyzed using the phasor approach^22, 23^, that provides a user-friendly graphical representation of fluorescence decays while allowing quantitative analysis. The Phasor analyzes were adapted to extract the FLT of the dyes in one image. The simplicity and specificity of phasor analysis for multiplexing different species provides a powerful research and discovery tool.

Our MC simulations show the correlation between the quantum efficiency of the fluorophore, its depth in the tissue, and our phasor-based analysis accuracy for extracting its FLT. We also investigated the multiplexing performance of our model as a function of depth in the tissue. We show that the accuracy of FLT calculations for the two fluorophores is significantly improved by applying a simple ‘cut-off” method to the resulting phasor analyzes. These new approaches were applied in our experiments and allowed discrimination between the two NIR dyes in an image behind tissue-like phantoms 1 cm thick.

## Methods

### Monte Carlo simulations

We present a new MC approach to study multiplexed FLI through tissue in the NIR regime. Our MC model is based on three main assumptions, that allowed us to simplify and shorten the computation time: (i) The model considers only the emitted fluorescent photons, and ignores the excitation photons (based on our recent publication^19^). (ii) The number of photons emitted into the tissue decreases according to a relaxation constant, that correlates with the lifetime of the fluorophore. (iii) Photons are collected by the detector at fixed time intervals, mimicking the gating mechanism in time-gated acquisition.

The fluorophore is located at a specific depth, *z*, in a semi-infinite tissue (infinite in dimensions x and y), with a refractive index *n*, an absorption coefficient *μ*_*a*_, a scattering coefficient *μ*_*s*_, and an anisotropic factor g (Figure 1(a)). We simulate the photons’ emission as an exponential decay, which correlates to a specific lifetime. Each point in Figure 1(b) represents a time point at which we sample the emitted photons, with time-intervals of 428 picoseconds (ps, correlated with the experimental time-gating parameters). The number of emitted photons in each gate step was determined using the exponential decay:

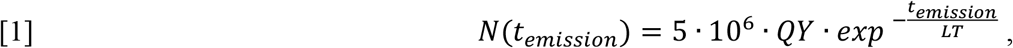

where *t_emi*ss*ion* = 428 ps. Each emitted photon propagated in the tissue, according to its scattering and absorption properties, for a nonspecific period of time, and then was either absorbed or exited the tissue surface into the air (*n* = 1). Photons reaching the detector were counted, within a given pixel (Figure 1(a)), while the total counted photons were multiplied by the SPAD array photon efficiency (PE, determined as 13%, corresponding to the SPAD array PE used in our experiments^21^). Once all photons emitted from a given time point on the decay curve have completed their propagation, the next emission occurs, described by the following point in Figure 1(b). All simulation parameters were chosen according to the characterization of the experiment (as described in the next paragraph), and are detailed in the caption of Figure 1.

**Figure 1.**
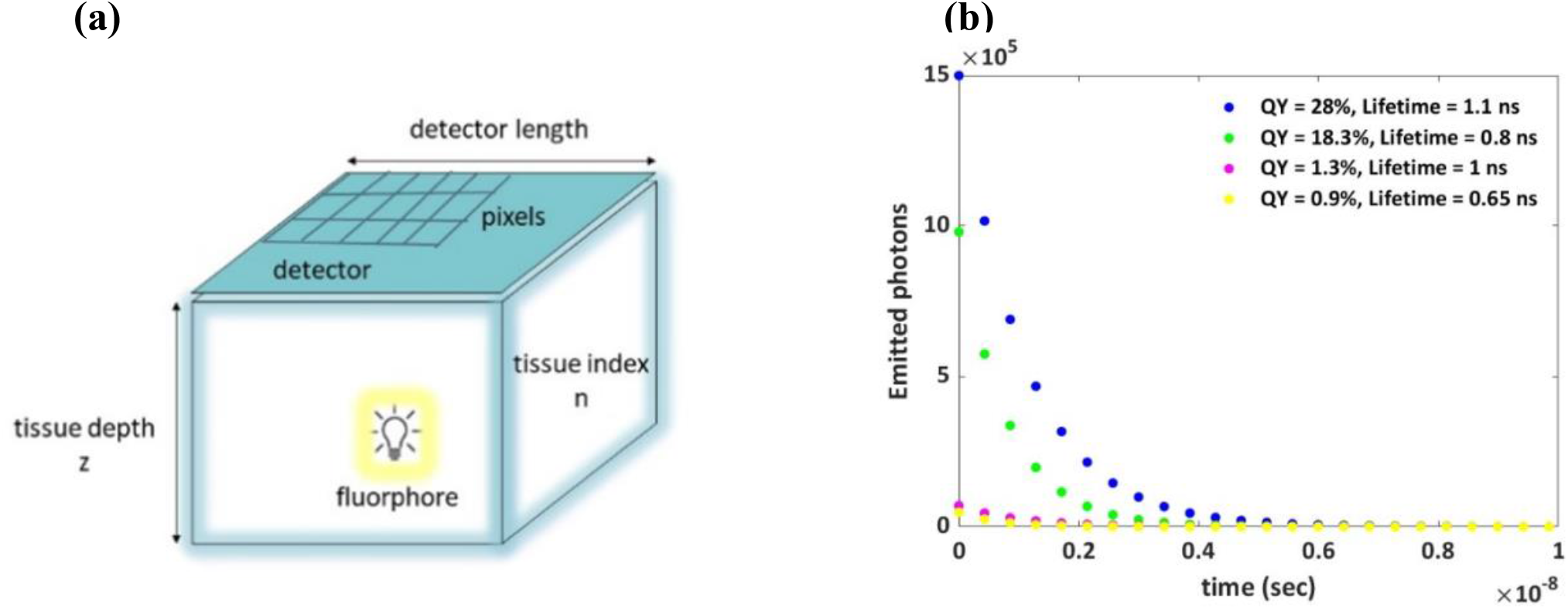
Simulation parameters. (a) A schematic description of the MC model simulating the propagation of fluorescent photons through a tissue of a given thickness z. Parameters used: Length of the detector =1 cm, number of pixels= 512*512 pixels, PE of the detector = 13%, refractive index n=1.4, scattering coefficient μ_s_=300 cm^-1^, absorption coefficient μ_a_=0.4 cm^-1^, anisotropy coefficient g=0.962^19^. (b) Emitted photons versus time, for different fluorophores with the following photophysical characterization: (i) Quantum yield (QY) = 28%, lifetime = 1.1 ns (blue dots). (ii) QY = 18.3%, lifetime = 0.8 ns (green dots). (iii) QY = 1.3% lifetime = 1 ns (magenta dots). (iv) QY = 0.9% lifetime = 0.65 ns (yellow dots). The time step between two sample points is g_s=428 ps, for 117 time samples (only the first 24 time points are shown in Figure 1(b)).

### Time-gated acquisition

The excitation source was a fiber-coupled pulsed laser with a wavelength of 779 nm, 20 MHz repetition rate, and a pulse width of ∼70 ps (VisIR-780, PicoQuant). Wide-field illumination was achieved with a 10x beam expander (GBE10-B, Thorlabs). The dye solutions were placed between two glass slides separated by a silicon insulator film (Merck, Israel). The emitted fluorescence was recorded with a NIR objective lens (5018-SW, Computar, TX) and imaged with the SPAD512S camera (PiImaging, Switzerland). The time-triggered in-pixel architecture enables time-resolved photon counting at a maximum rate of 97 kfps (1-bit frames). The photon detection efficiency of the detector was ∼13% at 800 nm with a fill factor of 10.5% and a dark count rate with a median value of 7.5 Hz/pixel. Overlapping gate images (G = 117, gate spacing g_s = 428 ps) were used, with a different gate image acquisition time (20 or 50 ms) in each experiment, to account for the different sample brightness (which decreased through the thickness of the phantom*s*).

### FLT phasor based analyzes

Phasor analyzes of the simulated photons, as well as of the experimental time-gated data, were performed as described in our previous paper^7^. Briefly, the uncorrected, uncalibrated phasor (g_i,j_, s_i,j_) of each pixel with coordinate (i, j) in the region of interest (ROI) in the fluorescence image was calculated as follows:

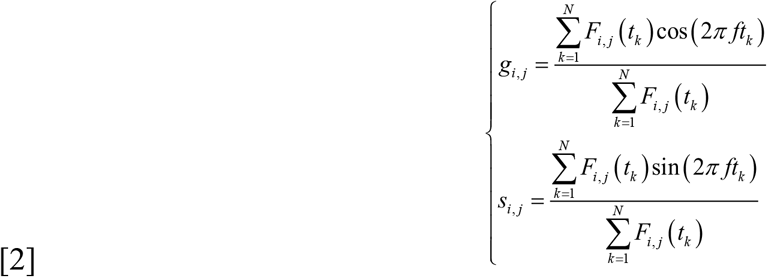

where f is the phasor harmonic (equal to the laser repetition rate = 1/T), k = 1 … N is the gate number and F_i,j_(t_k_) is the k^th^ gate image intensity value at pixel (i, j). The coordinates (*g*_*i,j*_, *s*_*i,j*_) can be plotted as a phasor plot with polar coordinates (*m*_*i,j*_, *φ*_*i,j*_) that are called modulation and phase, and are located on the universal cycle (UC)^22^:

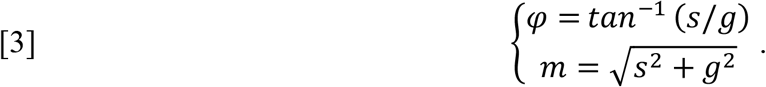

The phase lifetime, *τ*_*φ*_, was then computed using the following equation:

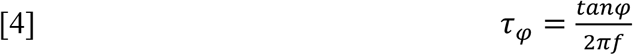

Phase lifetimes *τ*_*φ*_ were calculated for each pixel in the ROI, providing the data required for the histogram, which presents the number of counts for each *τ*_*φ*_. For each single dye, the phase lifetimes were calculated using the mean of the histogram, while for multiple dyes, the phase lifetimes were determined using the two highest columns of the histogram. In cases where the background noise was too high, or when the two adjacent fluorophores were very close to each other, we used a “cutoff” method in which the edge results were removed.

### Dyes solutions preparation and photophysical characterization

IRDye800 NHS (Li-Cor Biosciences, NE, U.S.A) and Indocyanine Green (ICG, Holland-Moran, Israel) were dissolved in Phosphate-Buffered Saline (PBS), to present a final concentration of 100 μM, then loaded onto glass slides spaced with silicon isolator sheets (Merck industries, Israel). The 3D plots of the excitation and emission spectra, for both dyes, was obtained using the Fluorolog-QuantaMaster system (Horiba Scientific, Jepan). Lifetime data were collected using the same system, equipped with a DeltaDiode-C1 controller (Horiba Scientific, Japan). Samples were excited with DeltaDiodes 730L and 830L pulsed lasers (Horiba Scientific, Japan), with a peak wavelength at 730nm/830nm +/-10nm and active temperature control with a repetition rate up to 100MHz. Emission was measured at wavelengths up to 900 nm. Fluorescence decay curves were analyzed using FelixFL decay analysis software (Horiba Scientific, Japan) based on a multiexponential model involving an iterative reconvolution process. The quality of the fits is assessed by the reduced χ2 value and a visual inspection of the distribution of the weighted residuals and their autocorrelation function.

The Strickler-Berg equation^24^ was employed to calculate the Dyes’ quantum yield (QY) from their fluorescence spectra, according to:

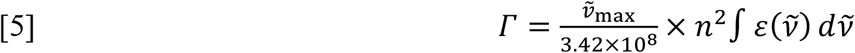

Where n is the refractive index of the medium,*v*_max_ is the position of the absorption maxima in wavenumbers [cm^-1^] and ϵ absorption coefficient. The resulted QY were 0.9% for the ICG dye and 3.3% for the IRDye800.

### Tissue-like phantoms of varying thicknesses

The tissue like phantoms were prepared of Intralipid and Ink, then solidified with agarose. The phantoms were prepared using India ink, as the absorbing component and Intralipid (IL) 20% (Lipofundin MCT/LCT 20%, B. Braun Melsungen AG, Germany), as the scattering component^21^. All phantoms contained the same ink concentration (0.008%) and IL (0.16%), resulting with an absorption coefficient of 0.04 mm^-1^ and a reduced scattering coefficient of 1.2 mm^-1^ (to mimic skin optical properties in the NIR regime)^25^. 1% agarose powder (SeaKem LE Agarose, Lonza, USA) was added to the solution to form a gel. The solutions were heated and mixed at a temperature of approximately 90 °C while the agarose powder was slowly added. The phantom solutions were stirred continuously to obtain high uniformity. The mixture was then poured into a large plastic plate and stored within water until used. For each experiment, a slightly oversized slice of phantom was cut and placed on a phantom holder comprised of a glass coverslip bottom taped to the 3D printed spacer of appropriate thickness (0.1, 0.3, 0.5, 0.7and 1 cm), then cut with a knife to achieve the desired thickness, and covered with another coverslip (see in Supporting information, Figure S7). These phantom slices are then slid on top of the sample (dyes within a gasket) within the 3D printed sample holder assembly.

## Results

### Simulations

Figure 2(a) shows a top-view of photons recorded by the detector throughout the simulated recording time, and for varying tissue thicknesses *z* (Figure 2(a) (i)-(vii), 0-1 cm). Figure 2(b) shows the average intensity <I> (counts) as function of tissue depth *z*, where the average intensity was calculated according to:

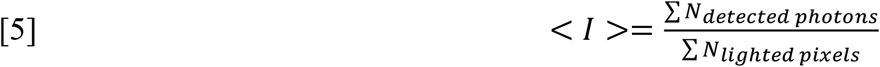

Clearly, intensity decreases as *z* increases, which is due to increased scattering and absorbing events. In addition, the intensity “spots” show higher scattering as the phantom depth increases.

**Figure 2.**
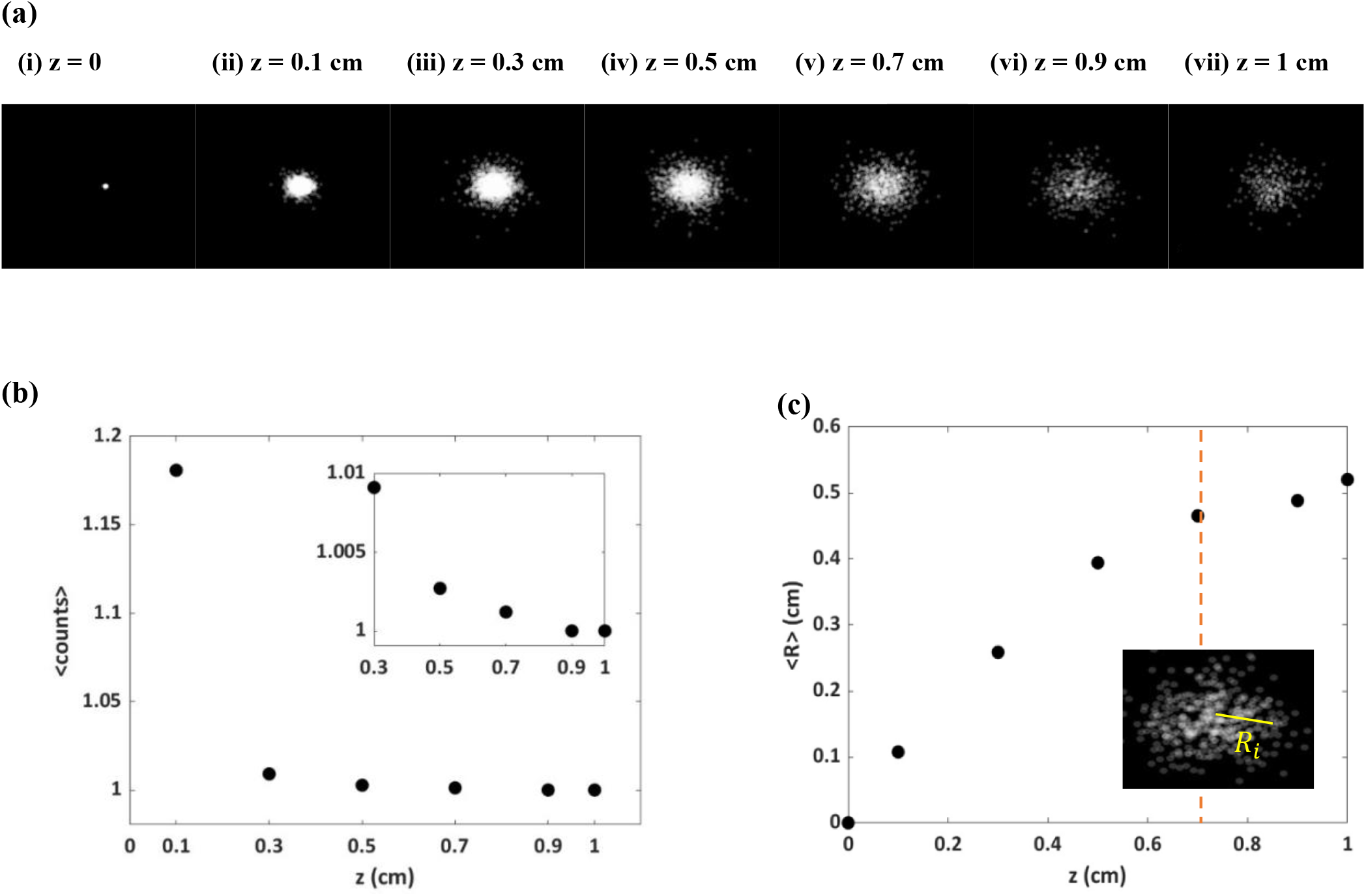
Simulated fluorescence intensity. **(a)** Snapshots of the captured photons in plan view for different tissue depths *z*: (i) 0 – (vii) 1 cm, using the MC simulation. **(b)** The average intensity <I> (counts) as function of z. <I> for *z* = 0 (no tissue) is 216 counts, and is not shown due to limited scale. Inset: <I> as function of z, for a depth range; 0.3-1 cm. **(c)** The average radius <R> as a function of z. Inset: a demonstration the of radius calculation *R*_*i*_ (yellow line) at z=0.7 cm.

Figure 2(c) shows the average radius R of the intensity “spot”, as function of tissue depth z. The radius for each detected photon (Figure 2(c), inset) was calculated from the center of the spot, and divided by the number of detected photons:

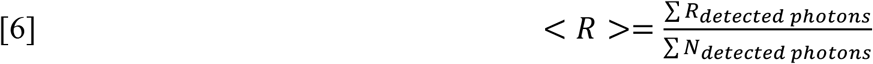

As expected, the average radius of the photons increases with the depth of the tissue, which is due to increased scattering events. Figure 2(c) shows an interesting behavior, where there is a strong dependance of < *R* > on small z values, that decreases with depth (dashed red line). This type of behavior, showing a lower dependance of fluorescence intensity on scattering at greater tissue depths, was also recently presented by Weber et.al, and explained by a novel diffusion theory of the paths of fluorescent photon in turbid media^19^. These overall results, which showed an expected, well-explained behavior, strengthened the accuracy of our MC model and paved the way for further FLT analyzes, as described below.

Once our model accuracy for wide field fluorescence imaging through a scattering medium was validated, we used phasor analyzes to extract the sample lifetimes from our simulated data. Figure 3 shows representative results for a phasor plot (3(a)) and lifetime analyzes (3(b)) for a single fluorophore with a lifetime = 0.62 ns (correlated with the experimental FLT of ICG, see in Figure 8) for an arbitrarily chosen depth: *z* = 0.3 *cm*. Phasor analyzes and histogram of lifetimes were calculated as descried in the *Methods* section above. Figure 3(b) shows the mean lifetime in the histogram, which is 0.65 ns, identical to the simulated lifetime of 0.65 ns. Similar results were obtained for all fluorophores shown in Figure 2(b) (data are available in the S*upporting information* section, Figure S1), as well as the results for various depths (*Supporting information*, Figure S2), while all the results are summarized in Figure 3(c). Despite the dispersed of the phasor points on the UC (rather than being centered as a single point, as expected for simulated data), the resulting lifetimes show a high correlation with the simulated lifetimes, for all tissue depths. Our model succeeded in predicting the paths of fluorescent photons in tissue, as well as to extract the correct fluorescence lifetimes for all samples.

**Figure 3.**
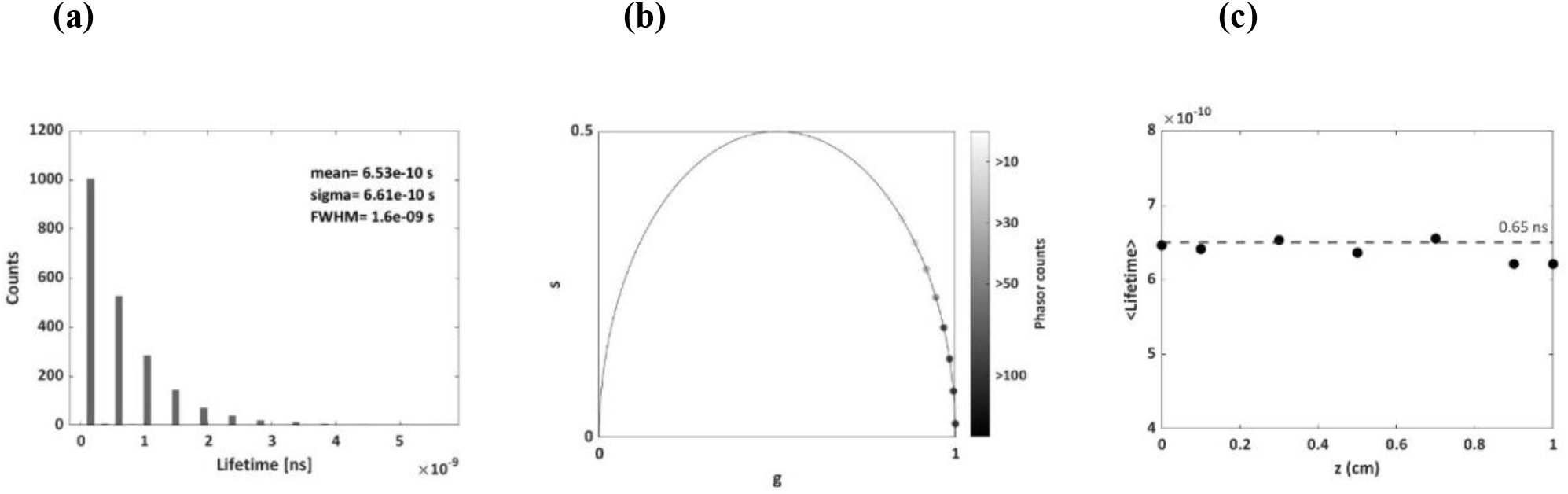
Representative phasor plot (a) and lifetime histogram (b) of a single fluorophore. The selected fluorophore has the following properties: QY = 0.9% lifetime = 0.65 ns, and is located at the depth of *z* = 0.3 *cm* within the simulated tissue (see Figure 2(a)). The mean lifetime extracted from the histogram perfectly matches the simulated FLT value. **(c)** Simulated FLTs extracted for all tissue depths, show a consistent FLT of 0.65 ± 0.01 · 10^−9^.

One of the most important issues to explore when considering FLI, is its performance in discriminating between two adjacent fluorophores, in one image, based on their lifetimes *τ*_1_, *τ*_2_. In this section, we describe simulated phasor-based analyzes for extracting lifetimes of two adjacent fluorophores with varying distances (Δ*x*), as a function of tissue thickness (*z*) and lifetimes separation (Δ*τ* = *τ*_2_ – *τ*_1_). Figure 4(a) shows a representative simulated intensity image of two adjacent fluorophores in plan view, located at a depth of *z* = 0.3 *cm* in the tissue. The optical properties of the fluorophores optical properties were determined from our experimental results (as described in the Methods section above): QY_1_ = 0.9%, *τ*_1_ = 0.62 ns (*F*_*1*_, yellow spot) and QY_2_ = 3.3%, *τ*_2_ = 1 ns (*F*_*2*_, magenta spot), and the vertical distance between the centers of the two fluorophores was Δx=2cm. The region of interest (ROI, the white rectangle) chosen for these analyzes is shown in Figure 4(a), inset, while Figure 4(b-i) shows the lifetime histograms and phasor analyzes for these ROI.

**Figure 4.**
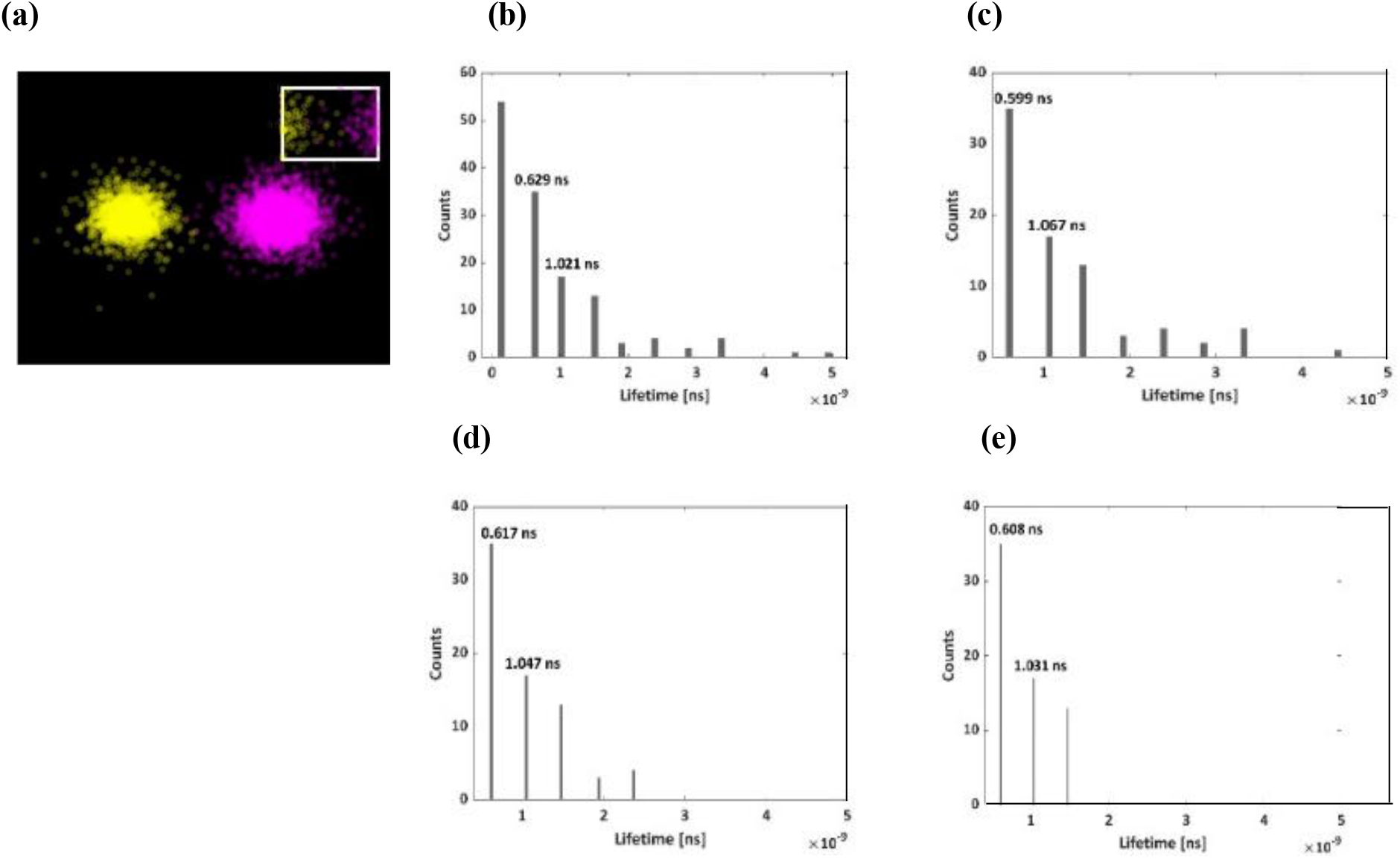
Simulated fluorescence intensity and phasor analyzes for two adjacent fluorophores placed at depth z = 0. 3 *c*m in the tissue, with a vertical distance Δ*x* = 2 *cm*. (a) The simulated intensity spots for the two fluorophores. The white square is the ROI, where the phasor analyzes were performed, and the inset is the magnification of this ROI. (b) Histogram of lifetimes for the two fluorophores in the ROI, resulting with lifetimes of 0.629 ns and 1.021 ns. (c-e) Histogram of lifetimes with a cutoff of 1% (c), 5% (d) and 10%(e) resulting with lifetimes of 0.599, 1.067 ns(c) 0.617, 1.047 ns (d), and 0.608, 1.031 ns (e). The cutoff was obtained by rejecting edge results from the total phasor values.

The highest column in Figure 4 (b) does not fit with any of the FLTs of the two fluorophores. The second column has a value of 0.63 ns, which is very close to *τ*_1_ (0.65 ns), and the third column shows 1.02 ns, also in high proximity to *τ*_2_ (1 ns). To make our results more precise and to avoid FLT values that do not correlate with the simulated FLTs, we performed an outlier exclusion, in which we set a threshold (1%, 5%, or 10%) for the resulting phasor values in the lifetime histograms. The lifetime histogram with a 1% cutoff value is shown in Figure 4 (c), the 5% cutoff value is shown in Figure 4 (d), and the 10% cutoff value is shown in Figure 4 (e). All cutoff results, even those with 1%, show better agreement with the known values and remove the first, high column. It is important to note that despite the analysis were performed on the pixels at the edge of the intensity spots (Figure 4 (a)), the results show a high correlation with the inserted parameters of the fluorophores, which is a clear confirmation of the performance of our new MC model in multiplexing different FLTs. Similar results were recorded for greater depth in tissue, of z=0.5 cm (see Figure 5 (a-c)) and z=1 cm (Figure 5 (d-f)), suggesting that lifetime extraction is not affected by the higher scattering events from the depth of tissue (a similar conclusion was also drawn in our previous article by Ankri et.al^7^). Encouraged by these FLT results, we then further challenged our new model: Figure 5 (g-j) shows the results for a smaller vertical distance, of Δx=1cm (measured by a vertical line connecting the centers of the intensity spots). Here, when the two intensity spots overlap, only the second and third columns show a relatively good fit to the FLTs of the fluorophores, leading to *τ*_1_=0.68 and *τ*_2_=0.96 ns. Rejecting an outlier with a 1% cutoff removes the edge result (first column in Figure 5(h)), leading to FLT values of 0.60 ns and 1.01 ns (Figure 5(i)), which correlate well with the simulated values. It is important to clarify that since only a few pixels overlapped (Figure 5(j), a magnification of Figure 5(g) ROI), the simple single-exponential analysis still allowed the correct FLT extraction.

**Figure 5:**
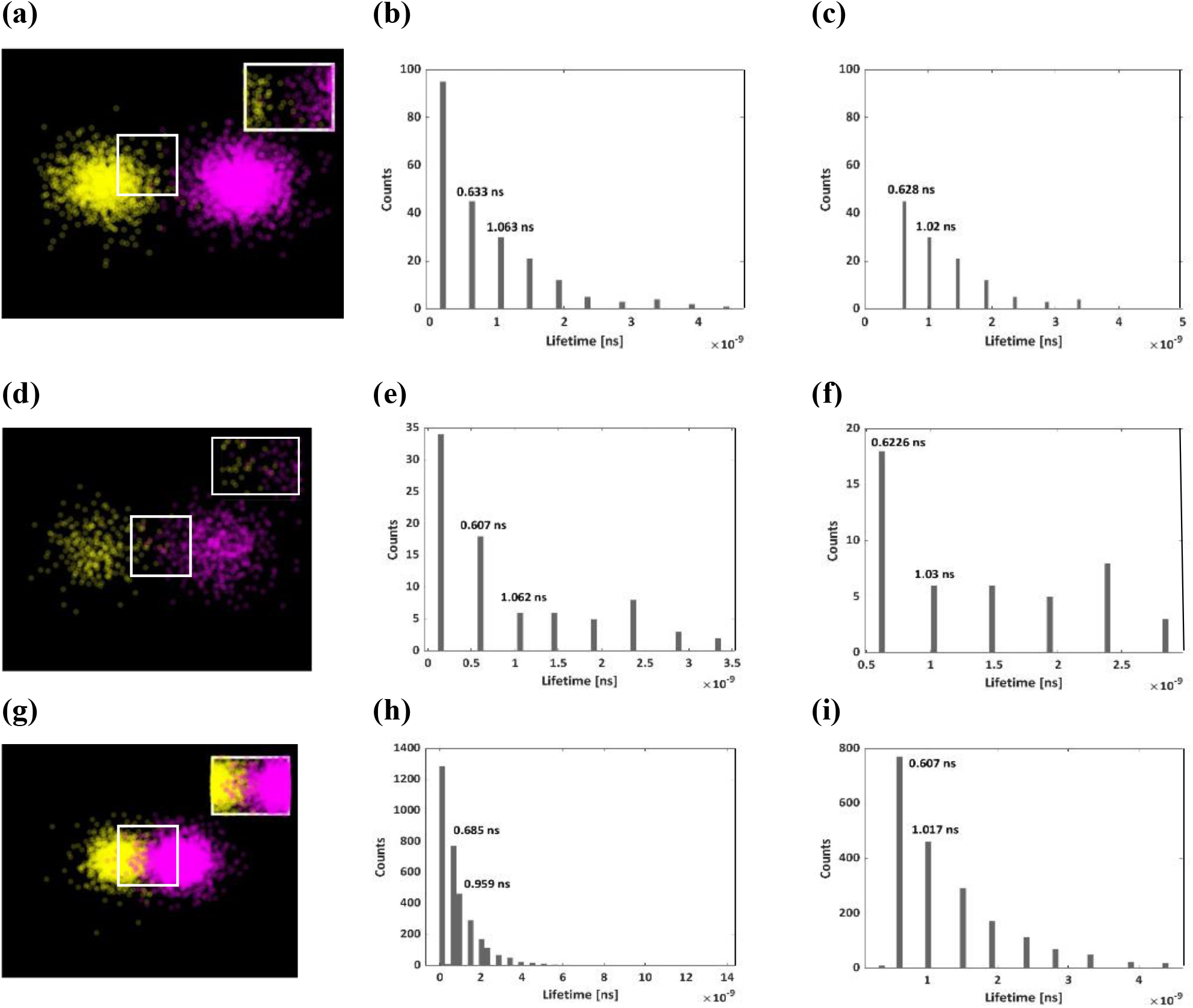
Simulated fluorescence intensity and phasor analyzes for two adjacent fluorophores. At all images, *F*_*1*_: yellow and *F*_*2*_: magenta. **(a-c)** z = 0.5 cm, Δ*x* = 2 *cm*. **(d-f)** z = 1 cm, Δ*x* = 2 *cm*. **(g-i)** z = 0.3 cm, Δ*x* = 1 *cm*.

Figure 6 summarizes the multiplexing results for the FLTs of the simulated samples (at the different depths *z* = 0, 0.1, 0.3, 0.5, 0.7, 0.9, 1 *cm*), without and with 1%, 5%, and 10% cutoff. The two fluorophores: *F*_*1*_ – yellow (Figure 6(a)) and *F*_*2*_ -magenta (Figure 6(b)) had a fixed spacing of Δ*x* = 2*cm* (the full data can be found in the *Supporting information* section, Figure S3). It appears that from a depth of 0.7 *cm* (dashed red line), the depth has a smaller effect on the resulting FLT, since the influence of the cutoff on the FLT is smaller. Interestingly, the depth *z* = 0.7 *cm*, is also the threshold depth that shows less dependance of fluorescence intensity on the scattering (Figure 2(c)). Figures 6(a) and 6(b) show a high correlation to the inserted FLT values, in all configurations, while higher cutoff values lead to more accurate results at greater depths.

**Figure 6:**
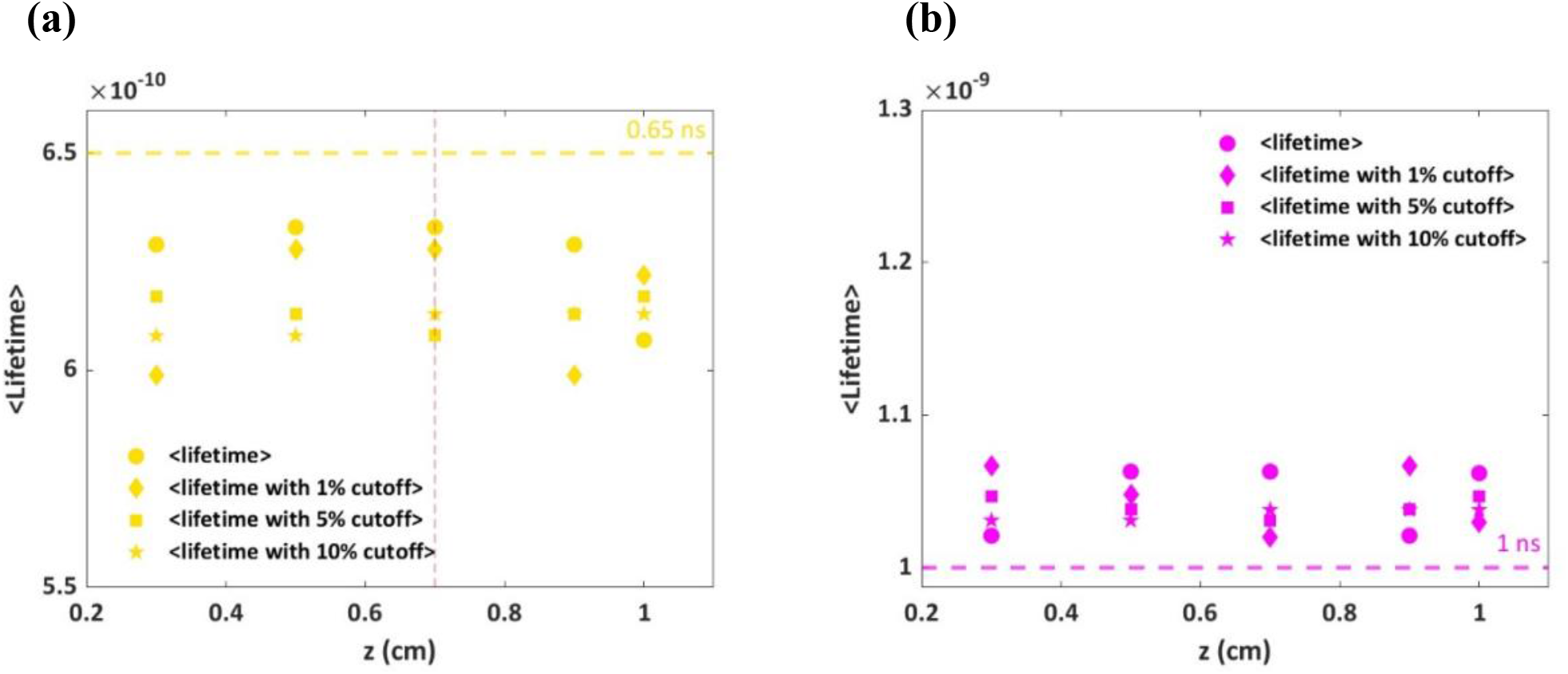
Lifetimes versus depth: without cutoff and with 1%, 5%, and 10% cutoff for two adjacent fluorophores with a fixed vertical distance Δ*x* = 2*cm*. **(a)** The fluorophore F_1_ **(b)** The fluorophore F_2_.

Identifying the limitations of a new model is an important step in its investigation. Therefore, studying the multiplexing capabilities of our new model for smaller distances was our next investigation. Figure 7 shows the extracted lifetimes for the vertical distances of the two adjacent fluorophores F1 and F2: Δ*x* = 1, 1.25, 1.5 1.75, 2 *cm*, for a fixed depth *z* = 0.3 *cm* (full data is available in *Supporting information*, Figure S4). The results show a high correlation with the inserted FLTs values, in all configurations.

**Figure 7:**
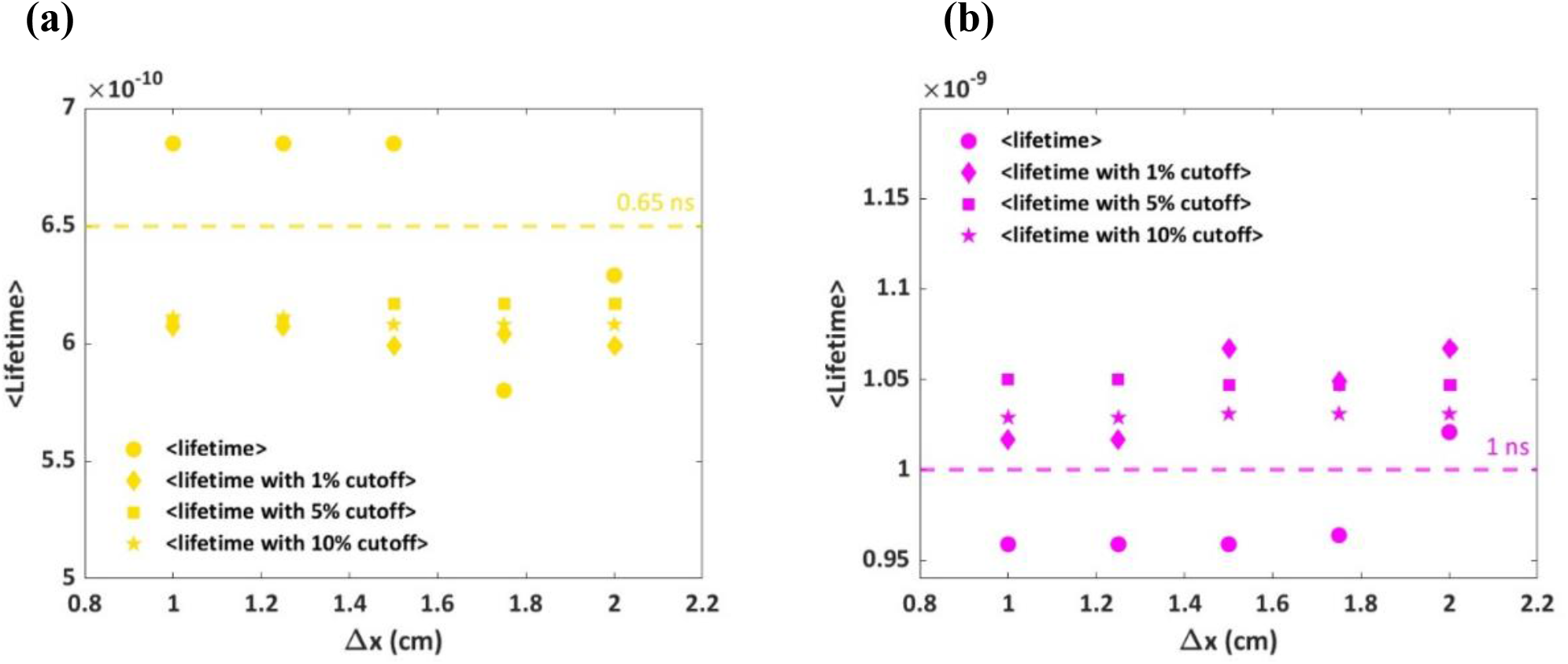
Lifetimes of two adjacent dyes for different vertical distances, Δ*x*. Different precents of cutoff were used: 1%, 5%, and 10% cutoff. **(a)** FLTs results for F_1_. **(b)** FLTs results for F_2_.

**Figure 8:**
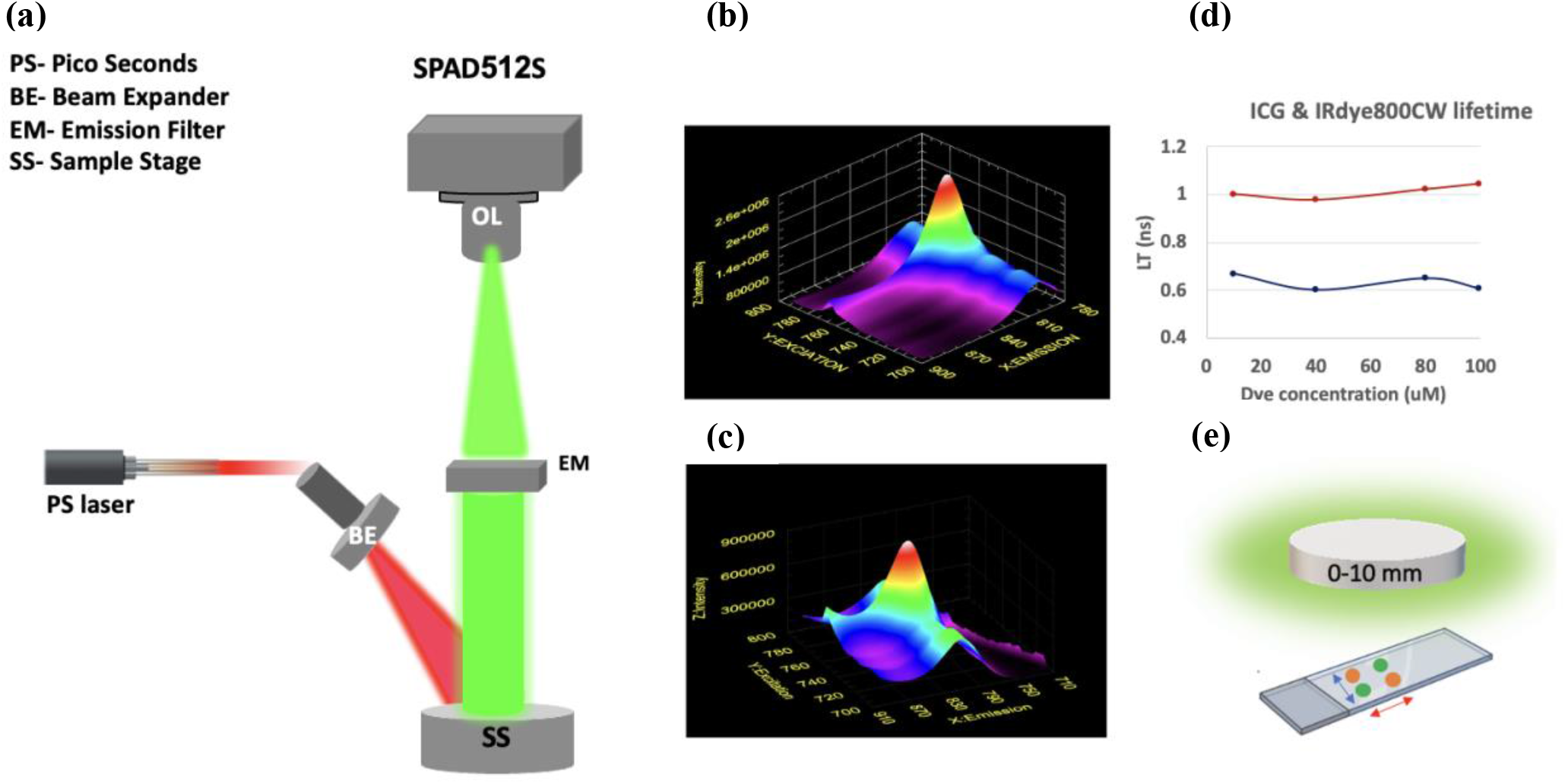
Parameters of the experiments: **(a)** The experimental setup used in the time-gated FLI: A 779 nm fiber coupled picoseconds (PS) laser illuminates the sample in a wide field mode through a X10 beam expander (BE). An objective lens (OL) collects fluorescence from the sample stage (SS) into the SPAD array through an emission filter (808 long pass filter, IDEX Health & Science, LLC, USA). **(b)** and **(c)** 3D emission and absorption spectra of the two fluorophores’ used in our experiments: IRDye800 (b) and ICG (c). The excitation peak for IRDye800 was at 770 nm and the emission peak was at 810 nm. For ICG, the excitation peak was at 785 nm and emission peak was at 820 nm. **(d)** FLT of the two dyes, measured for increasing concentrations of IRDye800 (red line) and ICG (blue line), using time correlated single photon counting (TCSPC). (e) The sample was prepared on a slide and spaced by a silicone gasket. The two dyes IRDye800 and ICG (orange and green) were applied to the slide at two distances of 0.5 and 1 cm, respectively (blue and red arrows, respectively). The phantom layers were applied to the slide at different thicknesses: 0, 0.3, 0.5, 0.7 and 1 cm..

### Experiments

Two commercial dyes, ICG and IRDye800, with measured lifetimes *τ*_1_ =0.62±0.03 and *τ*_2_ =1 ± 0.02, (Figure 8(d)) were placed at two specific distances, 0.5 and 1 cm, from each other. For each distance, phasor analyzes were performed, to extract *τ*_1_and *τ*_2_, and to study the scattering in the phasor cloud ^7^ as a function of Δx and the thickness of the phantom (the different Δx were achieved using silicon isolating sheets).

Figure 9(a) shows the intensity images of the two fluorophores (four different spots) imaged through phantoms’ of different thicknesses (0, 0.3, 0.5, 0.7 and 1 cm). Well seen, images become blurred as the thickness of the phantom increases, and the recognition of the four different dyes is impossible through the 1 cm thick phantom. Figure 9(b-d) shows the analyzes performed for ICG, the dye circled in red in Figure 9(a). Figure 9(b) shows the phasor “clouds” calculated for each pixel in the ROI. It is well seen, that the phasor cloud scatter increases with the thickness of the phantom, as was have previously reported by us^7^. From these phasor plots, we constructed the histograms describing the number of phasor counts for each FLTs (Figure 9(c)). Despite the increasing scatter in the FLTs, all samples had very similar mean FLTs, with a resulting average result of: 0.603 ± 0.022 nsec (Figure 9(d)). With the power of the microlensed SPAD512S, pixel-by-pixel background correction and phasor-based analyzes, the FLT of the dye could be extracted even behind a 1cm phantom.

**Figure 9:**
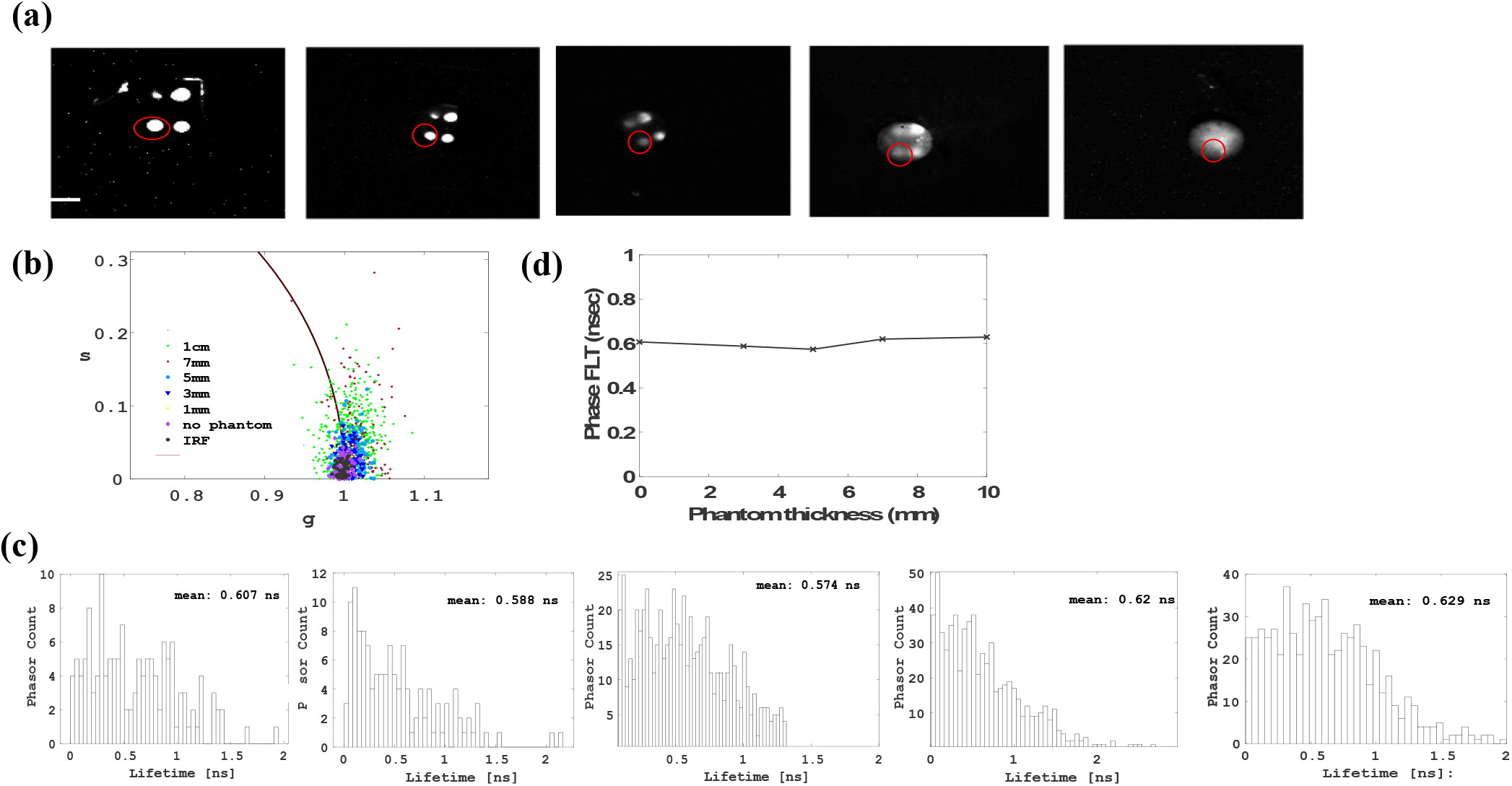
Extraction of ICG FLT behind an intralipid tissue-like phantoms with thicknesses varying from 0 to 1 cm. **(a)** The intensity images of the dyes for different phantom thicknesses. The scale bar is 12 mm. **(b)** Phasor “clouds” for the IRF and the different samples. The scatter in the “clouds” increases with the thickness of the phantom. **(c)** Histograms for the phasor counts versus the lifetimes calculated from each phasor point. **(d)** The mean FLT extracted for the dye behind the increasing thickness of the phantom.

Similar analyzes were performed for the IRDye800, and yielded an average FLT of 1.106 ± 0.058 nsec (Figure S8, *Supporting Information*), which is very similar to the FLT measured by our TCSPC setup (Figure 9(d)). These results indicate that phasor-based analyzes can perform FLI even through a 1 cm thick tissue like phantoms, by using the mean average calculations of the phasor cloud. For both dyes, the FLT increased slightly with the increasing thickness of the phantom, but the mean ‘phase FLTs’ were still very similar to the known FLT, 0.65 and 1 nsec the ICG and IRDye800, respectively, suggesting that despite the relatively small Δ*τ* between the two dyes, their multiplexing, using the same phasor analyzes, should be possible, as will be detailed hereinafter.

Once our method was established for a single-dye FLI behind tissue like phantoms, we used it for multiplexing purposes. Figure 10 shows the multiplexing performance of the phasor-based method for the two dyes behind a 0.5 cm thick phantom. We took the two dyes and calculated the phasor FLTs of their pixelated region (circled in red ROI). Because this region now also has a background (the region between the two bright spots), the resulting phasor shows long, spurious FLTs resulting from the BG in the ROI (Figure 10(b)), making it difficult to identify the FLTs of the two dyes. However, by choosing a time range in which the FLTs are expected to occur (i.e., 0-1.5 nsec, Figure 10(e)), we were able to extract the FLTs without a cutoff. Still, since our simulation results suggest a cutoff that should remove unwanted, long and/or short FLTs, Figure 10(c) shows the resulting phasor after a cutoff of 5%. With this cutoff, the phasor cloud is now much less scattered and shows a significant reduction in background noise. The phasor histogram shows two Gaussian distributions with two separate mean FLTs matching each FLT of the dye (Figure 10(g)). This result shows that by choosing the correct time range in the phasor histogram, with or without the cut-off method, we can easily multiply the FLTs of the two dyes in the intensity images.

**Figure 10:**
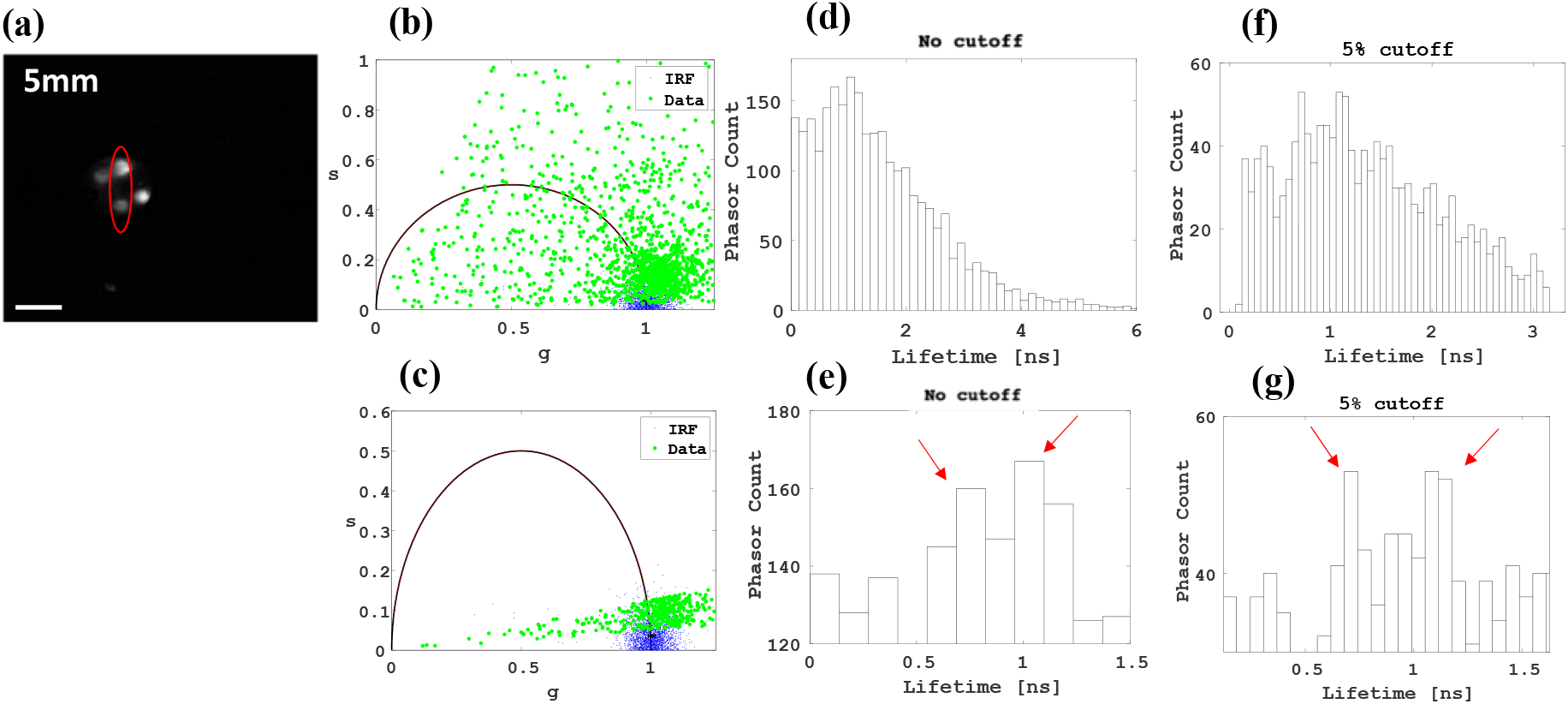
Multiplexing of ICG and IRDye800 behind a 0.5 cm intralipid tissue-like phantom. **(a)** The dyes circled in red (ROI) are the ICG and IRDye800. The scale bar is 15 mm. **(b)** and **(c)** Phasor scatter plot of the ROI without (b) and with a 5% cutoff. The phasor points of the data without cutoff are highly scattered due to background noise. Performing a 5% cutoff removes most of the BG noise. **(d-g)** Histograms for phasor counts versus lifetimes calculated from each phasor point for the different cutoffs performed. **(d**,**f)** Full phasor histogram. **(e**,**g)** Focusing on the 0-1.5 nsec region of interest where dye FLTs are expected to occur.

Similar analyzes were performed on data obtained during imaging through a 1-cm-thick phantom. Figure 11 shows the phase analyzes for the area circled in red, for different percentages of the cutoff: 5, 10, and 15%. The results indicate that by using a 1 cm thick phantom, the most accurate FLTs were obtained for a cutoff of 10%, with the highest columns representing the FLTs of 0.6 and 1.05, which correlate with the FLTs of ICG and IRDye800, respectively. With the single photon detection performance of SPAD512S and the phasor-based analyzes, the results suggest a simple multiplexing of two adjacent dyes after a single wavelength excitation. These time-gated, wide field FLI show that in the presence of a fluorescence signal sufficiently larger than the autofluorescence of the phantom, the fluorescence lifetime of the sample can be determined with an appropriate cutoff correction.

**Figure 11:**
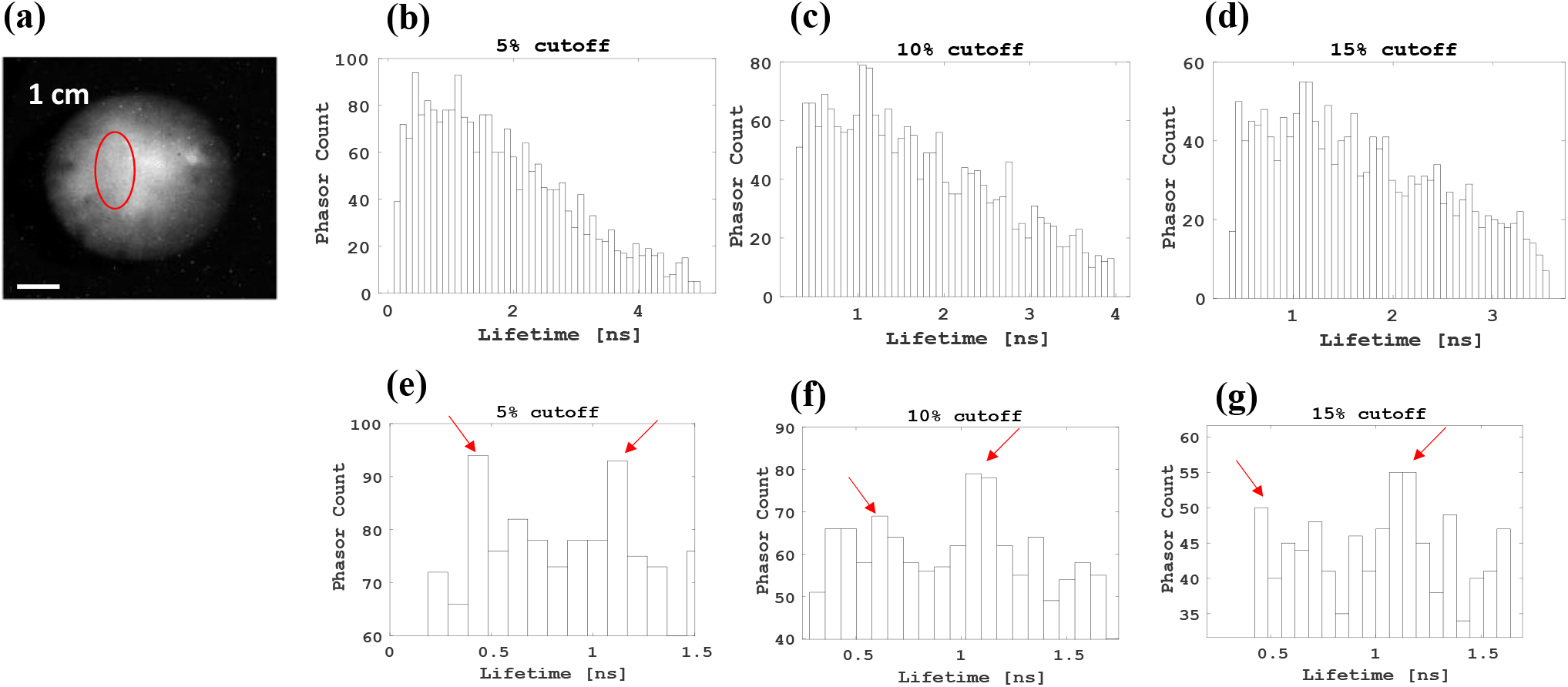
Multiplexing of ICG and IRDye800 behind a 1 cm thick intralipid tissue-like phantom. **(a)** The dyes circled in red (ROI) are the ICG and IRDye800 fluorescence. The scale is 15 mm. Phasor histograms of the circled ROI after a 5% (b), a 10% (c), and a 15% (d) cutoff. **(e-g)** Histograms of the phasor counts compared to the lifetimes calculated from each phasor point for the different cutoffs when focusing on the 0-1.5 ns time range of interest where the dye FLTs are expected to occur.

## Discussion

Fluorescence lifetime imaging (FLI) is widely utilized in life sciences, playing a critical role in biological and clinical diagnostics. For instance, in cancer research-FLI has been used for cancer cells detection^26, 27^, anti-cancer or chemotherapy drug delivery^28, 29^, and anti-cancer drug efficacy studies^30, 31^, enabled by distinguishing dyes with similar emission spectra by their FLTs, offering additional multiplexing capabilities. Due to the low autofluorescence and scattering of the tissue in the near infrared (NIR) range, as well as the emerging available fluorescence imaging techniques providing high sensitivity, noninvasive and quantitative readout, the NIR range of the electromagnetic spectrum is attractive for clinical fluorescence-based imaging methods. To this end, various instruments have been developed in the last decades for wide-field FLI required for in vivo measurements ^32-35^. One of the most widely used and significant instruments proposed for in vivo FLI is the timed SPAD array cameras ^36, 37^. SPAD cameras have been widely used in wide field FLI ^32, 38^. Thus, e.g., Ulka et. al. have proposed a large time-gated SPAD camera, SwissSPAD2, composed of 512×512 pixels^6^. Its FLI capabilities were demonstrated in the visible range, and the FLTs of organic dyes were extracted using phasor analysis. In another later work, they found that SwissSPAD2 provided consistent and high quality imaging of biologically relevant fluorescent samples stained with multiple dyes under various conditions^39^. They investigated the effect of gate length, number of gates, and signal intensity on the measured lifetime accuracy and precision of four commercially available fluorescent samples, as well as the ability of the sensor to quantify mixtures of two fluorescent dyes. They found that timed imaging with very long time gates did not hinder accurate determination of fluorescence lifetime as long as sufficient signal was recorded and the gate boundaries were well defined. Ankri *et. al*. ^7^ have investigated the effect of scatter on phasor-based FLT analysis. They have shown that for a single dye, the distribution of the phasor cloud (in pixels) increases with the broadening of the decay, which is due to scattering and decreasing fluorescence intensity from the depth of the tissue-like phantom. They also provided detection of several hundred thousand A459 cells expressing the fluorescent protein mCyRFP1 through highly scattering phantom layers, overcoming the scattering and autofluorescence of the phantom.

Multiplexing of different species in one image is easily enabled with the SPAD large-format cameras. Multiplexing FLI of different targets in one image is an important research niche with many potential applications ^40, 41^. Thus, e.g., Chen *et. al*. studied FRET (Förster Resonance Energy Transfer) kinetics in vitro and in vivo using a time-gated camera and a software tool (AlliGator) for phasor analysis^42^. The results for IgG binding kinetics, for both conventional matching and phase analysis, showed the same qualitative trends and confirmed the use of the phasor technique as an alternative tool for lifetime matching, showing FRET in vivo kinetics in bladder and liver. In another study, it was found that phasor analysis can be used to extract FLTs from mixtures of NIR dyes in in vitro and in vivo FRET^43^. Then, Smith *et. al*. extended this by reporting a macroscopic NIR in vivo measurement FLI-FRET (MFLI-FRET) using SwissSPAD245. They performed in vitro experiments ranging from monitoring environmental effects on fluorescence lifetime to quantifying FRET between dyes. They then successfully performed in vivo studies using FRET. Despite this extensive work on time-dependent phasor-based FLI in vitro and in vivo, none of this work has performed a comprehensive study of the multiplexing capabilities of this method as a function of dye quantum yield and tissue thickness, as we present in this work.

We first describe a new FLI simulation model based on some simple assumptions made by fast Monte Carlo simulations (MC) for clinically approved dyes with different lifetimes and quantum yields. The model describes the behavior of fluorophores at different depths within the scattering medium, followed by extraction of the FLT of the simulated dyes using the phasor approach. We demonstrate the FLI of two neighboring dyes at different spatial distances and depths, leading to a novel method that describes how to easily multiplex two targets in one image. We then confront our new method with experiments performed with a highly sensitive time-triggered camera SPAD. We show that our analysis allows multiplexing of FLI even within a 1 cm scattering phantom layer, overcoming the background autofluorescence signal and photon scattering events. The correlation between quantum efficiency and the number of counts within the lifetime histogram for two adjacent dyes needs further discussion. For the two parameters tested, depth in tissue and vertical distance between dyes, the highest number of counts in the lifetime histogram always results in the shorter lifetime, even when it is the FLT of the dye with the lowest quantum efficiency. This is surprising since it was expected that the dye with the lowest quantum efficiency would emit fewer photons, while the results contradict this assumption. Since this affected the strength of our method to extract the FLTs of the dyes, we did not investigate this further, but intend to do so in our future work. Our experimental results show that one can extract the FLT of a single dye from the FLT histogram using the mean of the histogram, even behind a 1 cm thick phantom. The “cut-off” method then made it possible to distinguish between two dyes in an image, according to their different FLTs in the histogram. Improving the image composition by increasing the intensity in the focal plane and using 3D imaging techniques, e.g. tomography, should improve the experimental results and allow greater depth of penetration into the tissue. To conclude, our MC FLI phasor-based model results, as well as our experimental SPAD-based wide field FLI, provide a deep study of phasor FLI for multiplexing purposes, providing a unique and simple method for the FLI of different targets in one frame.

## Acknowledgement

This work was supported by the Council for higher education, Israel (No. RA2100000248).

