## Supporting information for "Time-gated Fluorescence Lifetime Imaging in the Near Infrared Regime; A Comprehensive Study Toward In Vivo Imaging"

***Content:***

*Simulations:*

*Figure S1:* Phasor analyses and lifetime histogram for a single fluorophore with varying QYs.

*Figure S2:* Lifetime histograms and phasor analyses for $F_1$, for various tissues' depths

*Figure S3:* Fluorescence intensity and lifetimes histogram for two adjacent fluorophores $\Delta x = 2\ cm$ and varying depths.

*Figure S4:* Fluorescence intensities and lifetimes histograms for two adjacent fluorophores in various vertical distances $\Delta x$.

*Figure S5:* Fluorescence intensities and lifetime histograms for two adjacent fluorophores with a fixed vertical distance $\Delta x = 2\ cm$, and various depths within the tissue

*Figure S7*: Extracted lifetimes versus depth for varying precents of cutoff for two adjacent fluorophores with a fixed vertical distance $\Delta x = 2cm$.

*Experimental:*

*Figure S7:* The sliced tissue-like phantoms were used for the FLI experiments.

*Figure S8:* Extracting FLTs of IRDye800 imaged through intralipid tissue-like phantoms

*Supplementary information*

Simulation Results

Phasor analyses and lifetime histogram for <u>single fluorophore</u>, with varying QYs, that was placed in fixed depth inside the tissue, z = 0.3 cm, were calculated as detailed in the *Methods* section. (Results for $QY_1$ = 0.9%, $\tau_1$ = 0.65 ns are presented in Figure 3 in the paper).

**Phasor plot** **Lifetime histogram**

QY = 1.3%
Lifetime = 1 ns
(magenta in Fig. 1(b)
in the paper)

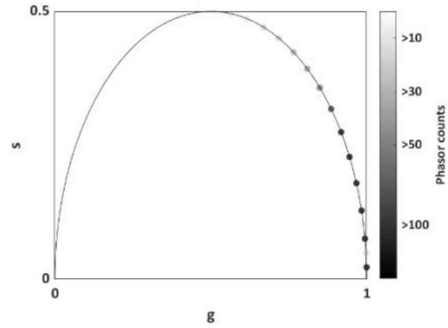


(a)

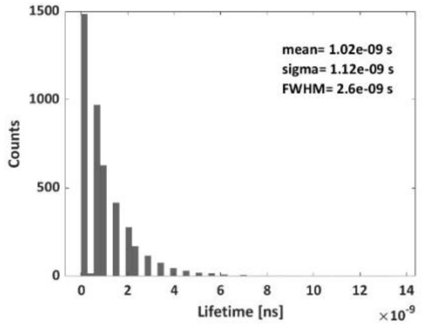


(b)

6

QY = 18.3
Lifetime = 0.8 ns
(green in Fig. 1(b)
in the paper))

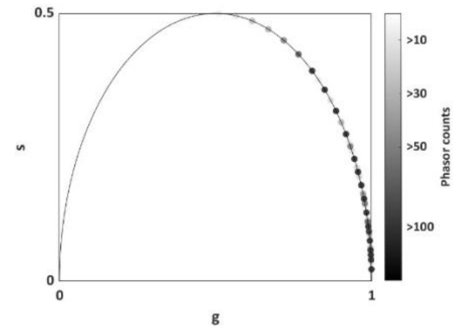


(c)

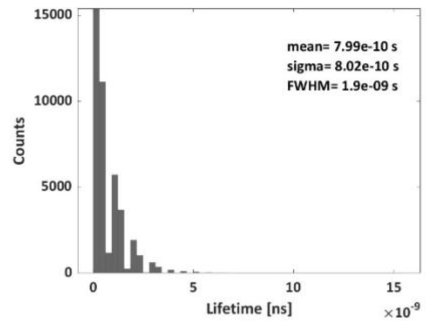


(d)

QY = 28%
Lifetime = 1.1 ns
(blue in Fig. 1(b)
in the paper))

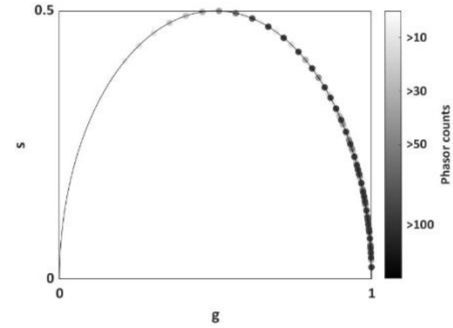


(e)

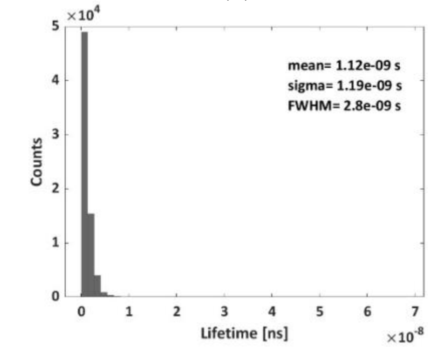


(f)

Figure S1. Phasor analyses and lifetime histogram for <u>single fluorophore</u> that was placed in fixed depth inside the tissue, z = 0.3 cm.

Phasor analyses and lifetime histogram for <u>single fluorophore</u> ($QY_1$ = 0.9%, $\tau_1$ = 0.65 ns) that was placed in various depths inside the tissue: (i) z = 0 – (vi) z = 1 cm, were calculated as detailed in the *Methods* section. (Results for z = 0.3 cm are presented in Figure 3 in the paper).

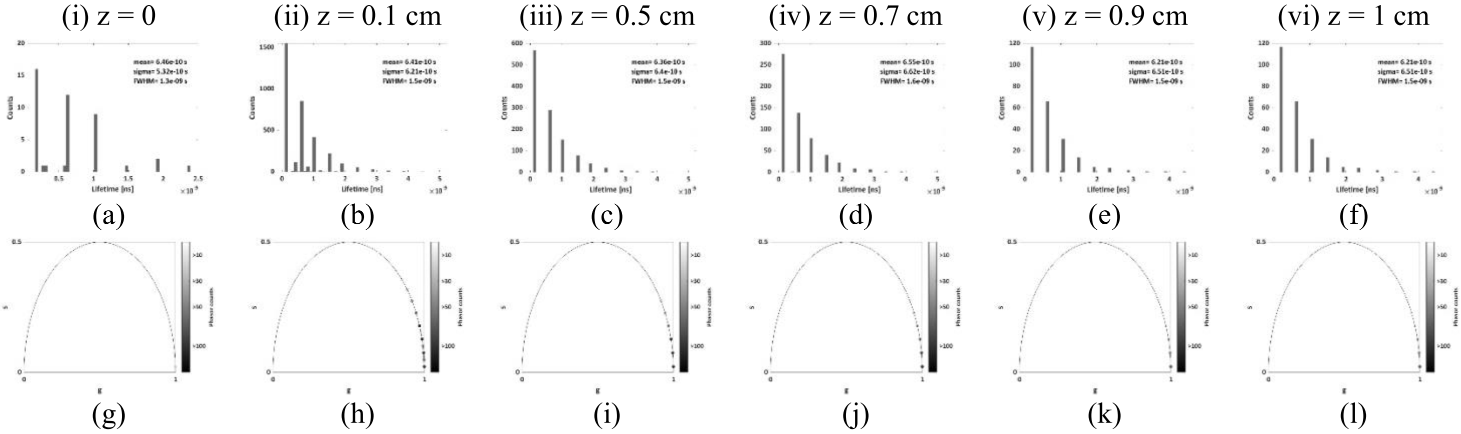


Figure S2. Lifetime histograms (upper row) and phasor analyses (bottom row) for $QY_1$ = 0.9%, $\tau_1$ = 0.65 ns for various depths inside the tissue: (i) z = 0 – (vi) z = 1 cm.

Simulated phasor-based analyses for lifetimes extraction of <u>two fluorophores:</u> $QY_1 = 0.9\%$, $\tau_1 = 0.62$ ns (yellow spot) and $QY_2 = 1.3\%$, $\tau_2 = 1$ ns (magenta spot), with fixed separation distances, $\Delta x = 2\ cm$, as function of the tissues' thickness, $z = 0{:}\ 1cm$ are presented in Figure S3.( Results for z = 0.3 cm are presented in Figure 4 in the paper). The ROI (white frame) for $z = 0.5 - 1$ cm is $1\ cm \cdot 1\ cm$ , whereas the ROI for $z = 0 - 0.1$ cm is $3\ cm \cdot 3\ cm$.

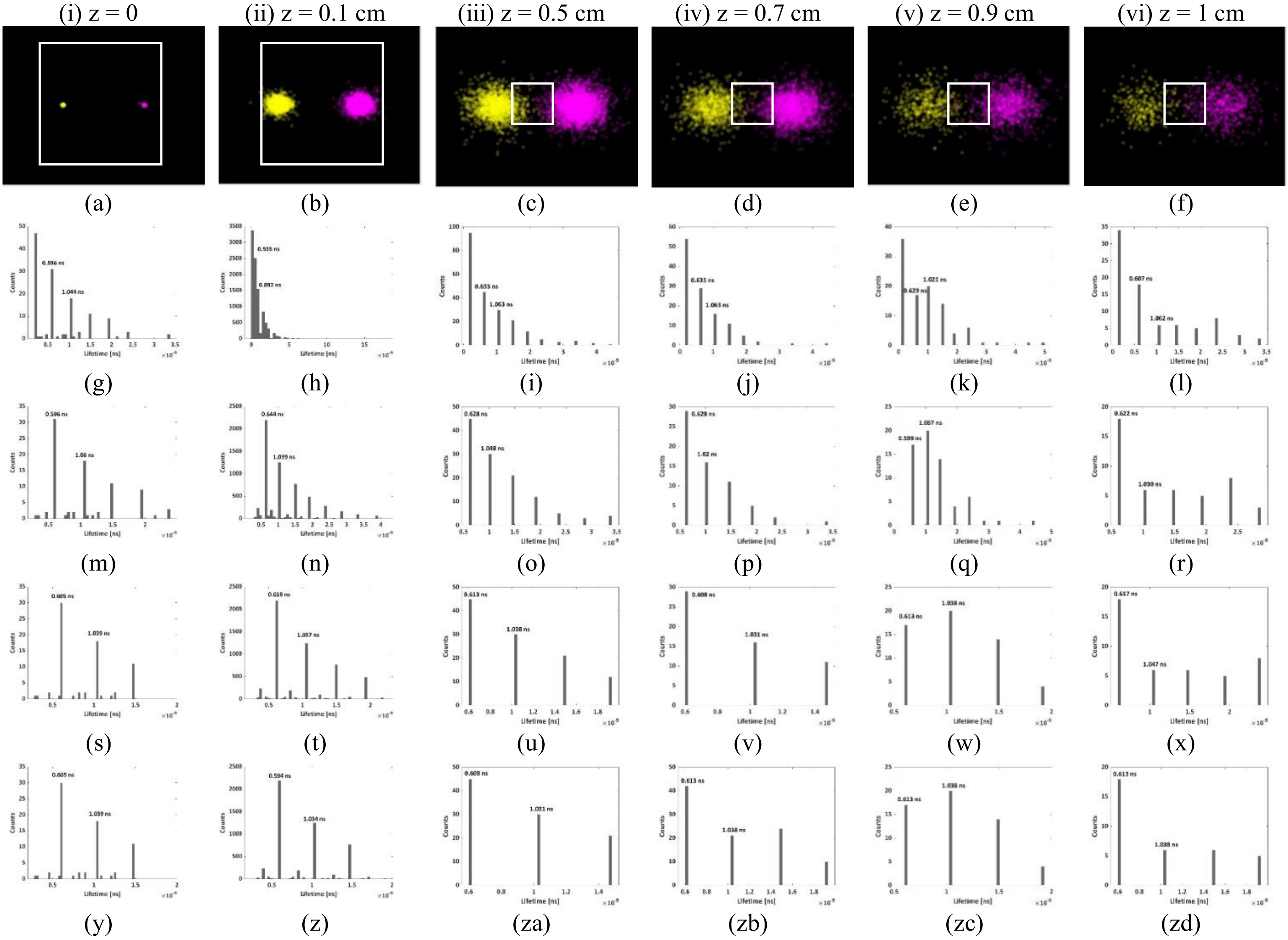


Figure S3. Simulated fluorescence intensity (first row) and lifetime histogram (second row) for two adjacent fluorophores with a fixed vertical distance $\Delta x = 2\ cm$ that were placed various depths inside the tissue: (i) z = 0 – (vi) z = 1 cm**.** Lifetime histograms with 1%, 5% and 10% cutoff are presented in the third, forth and fifth rows.

Simulated phasor-based analyses for lifetimes extraction of two fluorophores: $QY_1 = 0.9\%$, $\tau_1 = 0.62$ ns (yellow spot) and $QY_2 = 1.3\%$, $\tau_2 = 1$ ns (magenta spot), with fixed tissues' thickness, $z = 0.3\ cm$ as function of separation distances, $\Delta x = 1, 1.25, 1.5, 1.75\ cm$, is presented in Figure S4. The ROI (white frame) is $1\ cm \cdot 1\ cm$.

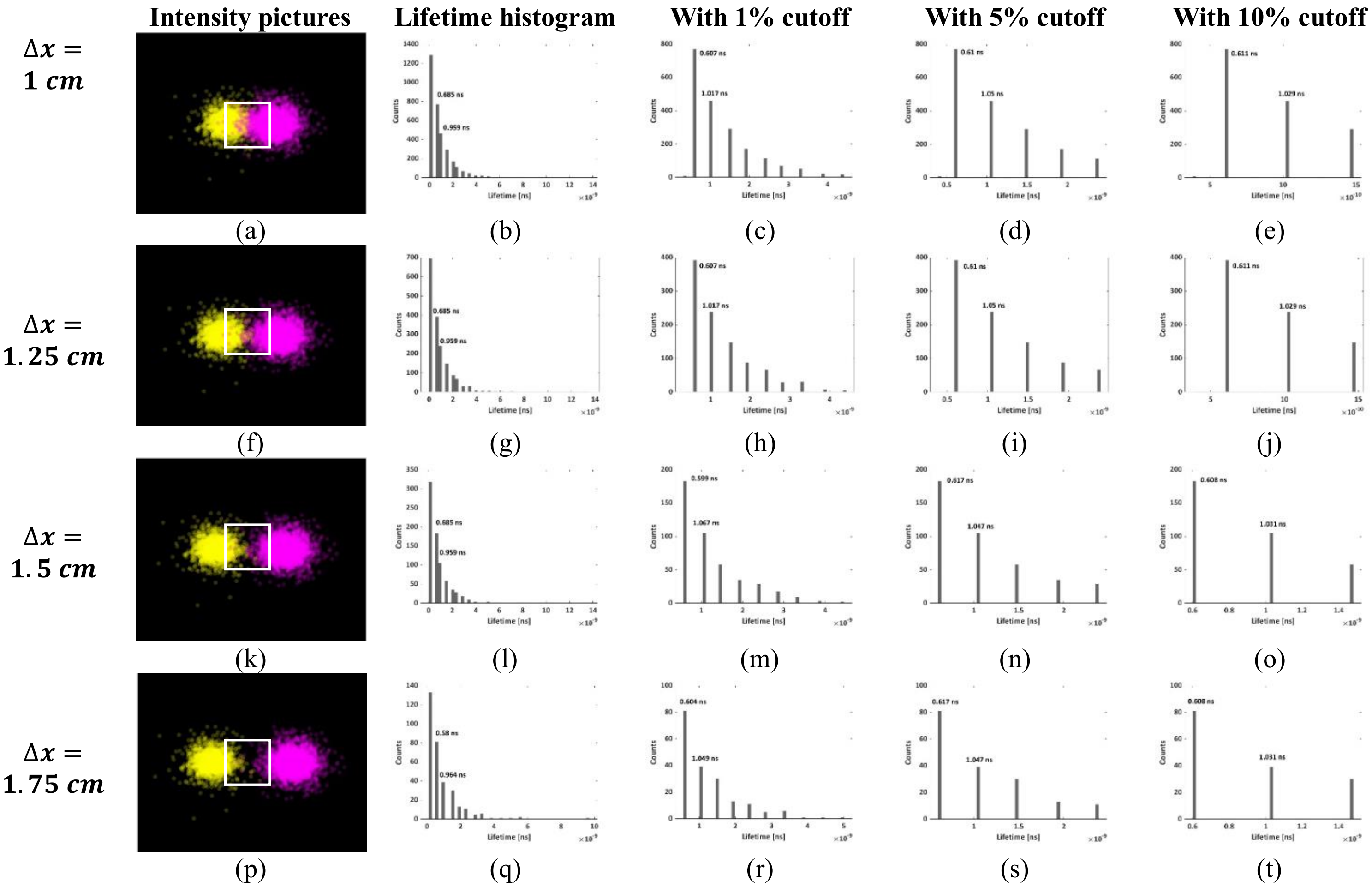


Figure S4. Simulated fluorescence intensities and lifetime histograms for two adjacent fluorophores with a fixed depth inside the tissue z = 0.3 cm that were placed in various vertical distance $\Delta x = 1, 1.25, 1.5, 1.75\ cm$**.** Lifetime histograms with 1%, 5% and 10% cutoff are presented as well.

Simulated phasor-based analyses for lifetimes extraction of two fluorophores: $QY_1$ = 18.3%, $\tau_1$ = 0.8 ns (green spot) and $QY_2$ = 28%, $\tau_2$ = 1.1 ns (blue spot), with fixed separation distances, $\Delta x = 2\ cm$, as function of the tissues' thickness, $z = 0{:}\,1cm$ are presented in Figure S5. The ROI (inset) is $1\ cm \cdot 1\ cm$.

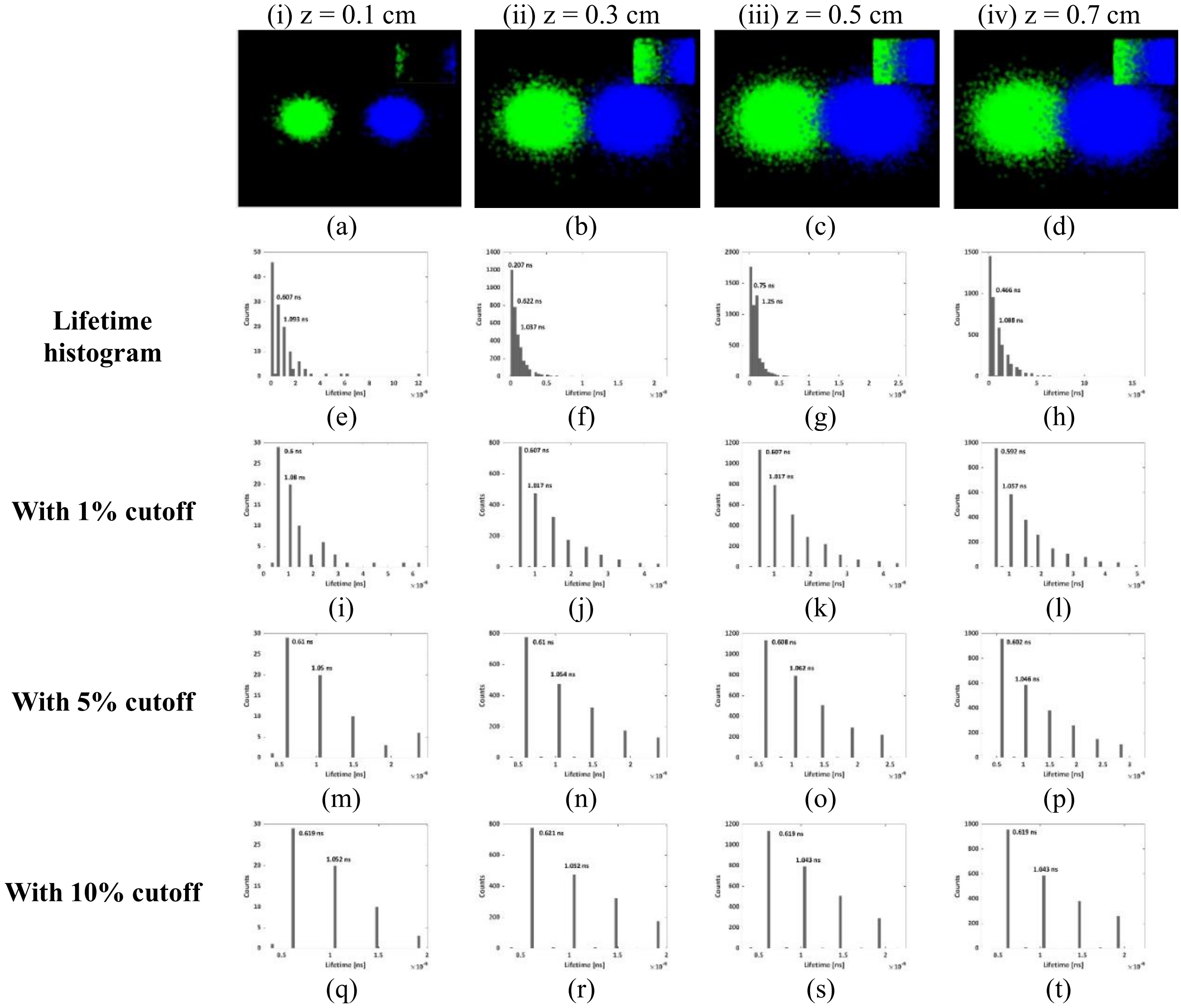


Figure S5. Simulated fluorescence intensities and lifetime histograms for two adjacent fluorophores with a fixed vertical distance $\Delta x = 2\ cm$, that were placed various depths inside the tissue: (i) z = 0.1 – (iv) z = 0.7 cm**.** Lifetime histograms with 1%, 5% and 10% cutoff are presented as well.

Figure S6 summarizes the multiplexing results for the FLTs of the simulated samples (at the different depths $z = 0.1, 0.3, 0.5, 0.7\ cm$), without and with 1%, 5%, and 10% cutoff. The two fluorophores: $F_3$ – green (Figure S6(a)) and $F_4$ -blue (Figure S6(b))) had a fixed vertical distance $\Delta x = 2cm$.

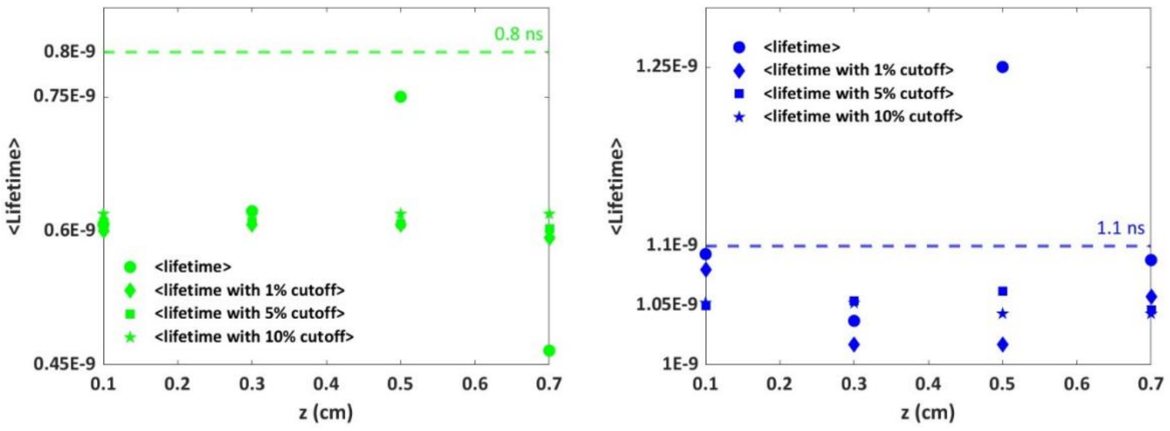


(a) (b)

Figure S6. Extracted lifetimes versus depth: without cutoff and with 1%, 5%, and 10% cutoff for two adjacent fluorophores with a fixed vertical distance $\Delta x = 2cm$. **(a)** The fluorophore $F_3$ **(b)** The fluorophore $F_4$.

**Experiments**

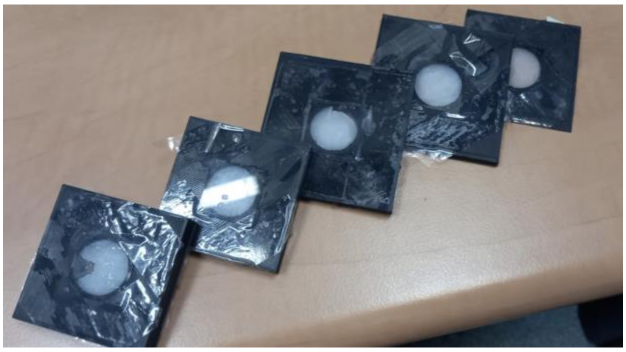

Figure S7: The sliced tissue-like phantoms were used for the FLI experiments. For each experiment, a slightly oversized slice of phantom was cut and placed on a phantom holder comprised of a glass coverslip bottom taped to the 3D printed spacer of appropriate thickness (0.1, 0.3, 0.5, 0.7 and 1 cm), then cut with a knife to achieve the desired thickness, and covered with another coverslip.

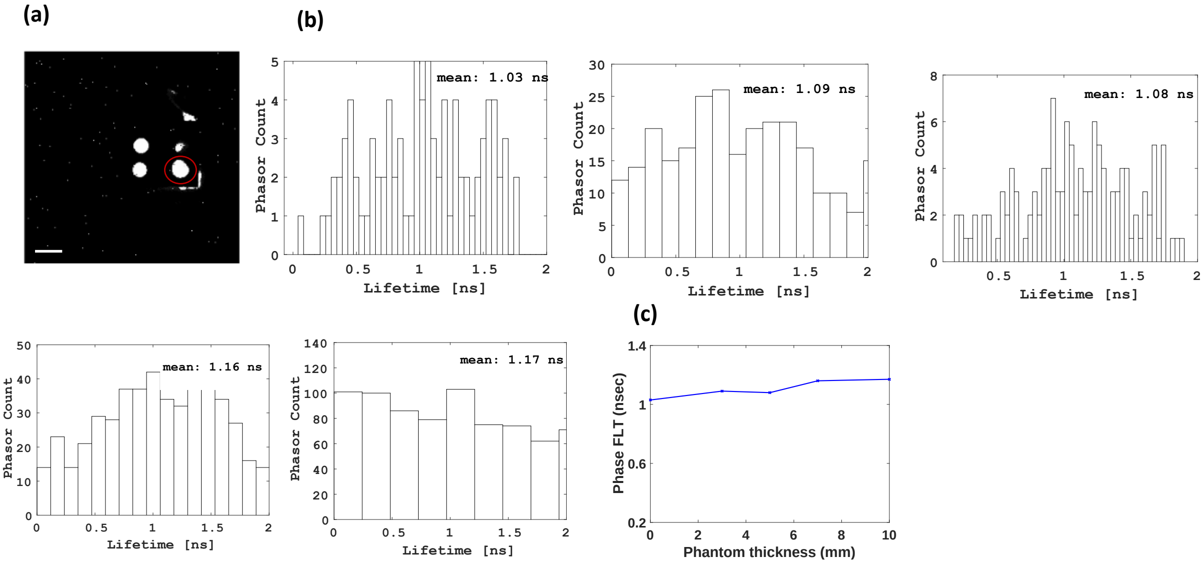


Figure S8: Extracting FLTs of IRDye800 imaged through intralipid tissue-like phantoms with thicknesses varying from 0 to 1cm. (a) The red circled dye is the IRDye800. Scale bar is 11mm. (b) Histograms for the phasor counts versus the lifetimes calculated from each phasor point. (d) The mean FLT for each phantom's thickness. (c) The mean FLT for each phantom's thickness. With the power of the microlensed SPAD512S, pixel-by-pixel background correction and phasor-based analyses, the FLT of the dye was extracted even behind a 1 cm phantom through the calculation of the mean of the phasor cloud.
